# F-box protein At1g08710 negatively regulates root length and imparts drought stress tolerance in *Arabidopsis thaliana*

**DOI:** 10.1101/2020.08.31.275271

**Authors:** Venkateswara Rao, Virupapuram Vijaybhaskar

## Abstract

Plants experience abiotic stresses throughout their life cycle and accordingly respond to tide over the unfavorable conditions. Drought or water deficit is one such condition to which plants respond in various ways including the ubiquitin proteasome system (UPS). Ubiquitin E3 ligases are a diverse family of protein complexes of which Skp1□Cullin□F□box (SCF) class mediate the ubiquitination and subsequent proteolytic turnover of proteins. F□box protein subunit plays crucial role in imparting specificity for selective degradation of target proteins. Here we report the function of *Arabidopsis* F-box protein *At1g08710* in drought stress adaptation. As F-box protein is a constituent of SCF complex, here it is shown interacting with ASK1 and Cullin1. F-box protein localizes to both nucleus and membrane. F-box gene transcript accumulates highly in root and is altered in response to drought stress conditions. F-box protein interacts with a transcriptional co-activator protein ADA2b. F-box mutant plants displayed better growth under drought stress conditions compared to the wild type with a reduced accumulation of H_2_O_2_ and malondialdehyde (MDA). Drought responsive genes *RD29A, RD22, ABI3* expression is also induced in F-box mutant plants. These results indicate F-box protein At1g08710 plays a role in drought stress adaptation in *Arabidopsis thaliana*.

**Highlights:**

- F-box gene *At1g08710* encodes a nuclear, membrane localized protein.
- F-box protein At1g08710 interacts with transcriptional co-activator protein ADA2b.
- F-box protein imparts drought stress tolerance by modulating H_2_O_2_ and MDA content.
- Drought response genes *RD29A, RD22, ABI3* expression is induced in F-box mutant plants.

## 1. Introduction

Plants being sessile are invariably prone to various environmental stress conditions throughout their life cycle. Abiotic stress conditions such as drought, heat, osmotic and cold adversely affect plant growth and crop productivity. Among these stress conditions, drought severely affects rain fed agriculture and plant productivity. Among various plant responses plant hormone Abscisic acid (ABA) plays an important role in drought stress adaptation. Phytohormone ABA induces the stomatal closure to prevent the water loss under drought stress condition (Finkelstein et al., 2002; Raghavendra et al., 2010). Drought stress induces the production of ABA, which in turn activates the expression of many drought stress responsive genes such as Responsive to Dehydration 29A/B (RD29A/B), Cold-Regulated Protein 47 (COR47), Kinase1(KIN1) and Responsive to ABA 18 (RAb18) (Abe et al., 2003; Kurkela and Borg-Franck, 1992; Lang and Palva, 1992). In addition to this, various dehydration proteins are also accumulated in plants in response to drought stress such as antioxidants, chaperones and late embryogenesis abundant (LEA) proteins (Goyal et al. 2005; Mittler 2002; Sun et al. 2002).

Recent reports suggest ubiquitin proteasome system (UPS) plays a critical role in plant response to abiotic stress conditions. Ubiquitin ligase E3s are key enzymes of this pathway which provide substrate specificity. Role of E3 SCF ligase complex proteins in plant adaptation to various abiotic stress conditions has been revealed recently (Lyzenga et al., 2012). F-box protein is an integral member of E3 ligase SCF complex comprising F-box protein, Cullin, Rbx1 and SKP1 proteins. Each component of this multi-subunit enzyme plays an important and distinct role. F-box proteins contain their characteristic F-box domain consisting 40-50 amino acid residues, approximately. In F-box proteins, degenerate F-box domain is present on the N-terminus region whereas different variable domains are positioned on C-terminus region such as WD40, Kelch repeat, Leucine rich repeat (LRR), Tetratricopeptide repeat (TRR) which are required for the interaction with target proteins. F-box proteins can interact with one or multiple SKP proteins and thus involved in the formation of functional SCF complexes. However, some F-box proteins are known to function independent of SCF complex (Gagne et al., 2002). Regulation of various biological processes such as circadian clock, photo morphogenesis, self-incompatibility, hormone signaling and floral organ identity are determined by F-box proteins. UFO is an F-box protein found to be involved in floral organ development (Durfee et al., 2003). LKP1, ZTL, LKP2 and FKL are involved in circadian rhythms and photomorphogenesis (Somers et al., 2000). TIR1 F-box protein involved in auxin signaling and identified as auxin receptor (Dharmasiri et al., 2005). SLY1, SNE F-box proteins regulates gibberellic acid signaling (Ariizumi et al., 2010). F-box protein COI1 has been implicated in jasmonic acid signaling (Katsir et al., 2008).

F-box proteins participate in various biological functions by interacting with target proteins, which often leads to the degradation of such proteins through ubiquitin mediated 26S proteasome pathway. Recent studies have revealed involvement of F-box proteins in stress adaptation. *Arabidopsis thaliana* RCAR3 INTERACTING F-BOX PROTEIN 1 (RIFP1) is involved in ABA signaling. RIFP1 interacts with regulatory components of ABA receptor 3 (RCAR3). Mutant plants of *rifp 1* exhibited tolerance to drought stress. Whereas, RIFP1 overexpressing plants are insensitive to ABA treatment. The rifp1 mutant plants displayed ABA mediated inhibition of seed germination. ABA responsive *AB18, RD29A, RD29B, ABF3* genes expression is induced in *rifp1* mutant plants (Li et al., 2016). F-box protein Drought Tolerance Repressor 1 (DOR1) is involved in ABA mediated drought stress tolerance. Mutation in the *DOR1* gene induces stomatal closure under water deficit conditions, leading to drought stress tolerance. Whereas, plants overexpressing *DOR1* are sensitive under drought stress conditions (Zhang et al., 2008). SCF E3 ligase complex subunit, Phloem protein 2-B11 (AtPP2-B11) negatively regulates drought stress. The expression of AtPP2-B11 gene was significantly induced under drought stress treatment. AtPP2-B11 interacts with AtLEA14 protein reducing its stability under drought stress conditions. AtPP2-B11 overexpressing plants are hypersensitive to drought stress and displayed altered expression of stress inducible genes such as *COR15a, COR47, ERD10, KIN1, RAB18*, and *RD22* (Li et al., 2014). ABA-responsive FBA domain-containing protein 1 (AFBA1) is a positive regulator of ABA response and involved in drought stress tolerance. *AFBA1* overexpressing plants exhibited tolerance to drought stress and displayed ABA mediated rapid closure of stomata. Whereas, *afba1* mutant plants are sensitive to drought stress and stomatal movement is ABA-insensitive (Kim et al., 2017). F-box protein More axillary growth 2 (MAX2) has been shown to be involved in drought stress. Arabidopsis *max2* mutant plants are hypersensitive to drought stress compared to the wildtype plants. ABA mediated stomatal closure is impaired in *max2* mutant plants (Bu et al., 2014). So, the genetic and biochemical evidence in Arabidopsis thaliana suggests that post-translational turnover of proteins mediated by SCF complexes is important for the regulation of diverse developmental and environmental pathways of which F-box proteins specify target proteins for degradation through UPS.

In the present study, *Arabidopsis thaliana* F-box protein encoding gene *At1g08710* was cloned and functionally characterized. Initial transcript analysis revealed its higher expression in root followed by other organs of mature plant. F-box gene transcript is altered in plants treated with mannitol and Abscisic acid. Interaction study revealed F-box protein’s interaction with transcriptional co-activator protein ADA2b. F-box gene mutant plants are able to survive better under drought stress conditions compared to the wild type plants. Expression of drought responsive genes *RD29A, RD22, ABI3* are induced in F-box mutant plants indicating F-box protein At1g08710 involvement in drought stress adaptation.

## 2. Materials and Methods

### 2.1 Plant material and growth conditions

*Arabidopsis thaliana* ecotype Col-0 seeds were used in all experiments. All plants were grown in plant growth chamber under the controlled conditions of 16/8 h light/dark cycle, 22 ± 2 °C and a light intensity of 100 μm m^−2^ s^−1^.

### 2.2 Isolation and molecular cloning of F-box protein At1g08710

To clone the F-box gene coding sequence (CDS) from Arabidopsis, total RNA was isolated from roots using Tri reagent (Sigma, USA) and cDNA was prepared. Primers were designed on the basis of reported cDNA sequence in TAIR (*At1g08710*) and PCR was done using Phusion DNA polymerase. The amplified PCR product was cloned in to pENTD-TOPO vector.

### 2.3 RNA extraction and quantitative real time PCR

Total RNA was isolated from different organs of *Arabidopsis thaliana* mature plant using TRI reagent (Sigma). ABI step one real time PCR was used for the quantitative transcript analysis. Expression of target gene was normalized with the expression of endogenous control *18S rRNA*. 20μl Real-time PCR reaction contains 10μl of Power SYBR green PCR master mix (Applied Biosystems), 2μl (1:20) diluted cDNA product, 20 pmole each of forward and reverse primer. For each primer set no template control was included as negative control. ΔΔCT method (Applied Biosystems) was used to calculate relative expression of each gene. All reactions were performed in triplicate with at least three biological replicates.

### 2.4 Yeast two hybrid experiment

#### Cloning of At1g08710 in pGBKT7BD vector

F-box gene *At1g08710* CDS was amplified from cDNA using gene specific primers (Supplementary data) and cloned in pENTD-TOPO vector. *At1g08710* was further subcloned in yeast two hybrid vector pDEST-GBKT7 using Gateway cloning technology (Invitrogen, CA). Cloning was confirmed by colony PCR and sequencing.

#### Yeast two hybrid (Y2H) experiments

Protein interaction analysis was performed by Yeast two hybrid system (Clonetech). Y2H Gold, Y187 yeast strains were used to carry out the experiments. F-box gene *At1g08710* was subcloned in to bait vector pDEST-GBKT7 and transformed into Y2H Gold strain using EZ-Yeast Transformation Kit (MP Biomedicals). This construct was mated with Arabidopsis normalized mate & plate library, which is high-complexity cDNA library cloned into a pGADT7AD vector and transformed into yeast strain Y187. These transformed cells were initially selected on SD/-Leu/-Trp (DDO) media. Interactions were confirmed by further screening on SD/-Leu/-Trp/-His/-Ade/X-α-gal/Aureobasidin (QDO/X/A) media. Cells which could grow on QDO/X/A and produce blue color were considered positive for interaction. pGADT7-T and pGBKT7-Lam vectors used as negative control and pGADT7-T and pGBKT7-53 used as positive control. EZ Yeast plasmid preparation kit (G-Bioscience) was used to isolate the plasmids from these colonies. These plasmids were further sequenced by using GAL4AD vector specific primers to identify the interacting proteins.

#### Yeast 2 hybrid one to one interaction

To further confirm the interaction of F-box protein At1g08710 with transcriptional co-activator protein ALTERATION/DEFICIENCY IN ACTIVATION 2B (ADA2b), *At1g08710* was cloned in GAL4 DNA-binding domain expression vector pDEST-GBKT7 and *ADA2b* was cloned in GAL4 activation domain expression vector pDEST-GADT7. These two constructs were transformed to Y2H gold strain and grown on SD/-Leu/-Trp/-His/-Ade/X-α-gal/Aureobasidin (QDO/X/A) agar media. Cells which are able to grow and produce blue color on QDO/X/A were considered positive for interaction. Positive and negative controls were also used in this experiment as supplied by the kit.

### 2.5 Bimolecular fluorescence complementation (BiFC) assay

Vectors pSAT4-DEST-N (1–174 N-YFP) and pSAT5-DEST-C (175-END C-YFP) were used for BiFC. F-box gene *At1g08710* was cloned in pENTD-TOPO vector and transferred to pSAT4-DEST-N (1–174 N-YFP) and *ADA2b* was cloned in pENTD-TOPO vector and transferred to pSAT5-DEST-C (175-END C-YFP) vector using Gateway cloning technology (Invitrogen, CA). BiFC was done in onion epidermal cells using PDS-1000 Helios Gene Gun (Bio-Rad) according to the manufacturer guidelines. These onion peels were kept on Murashige and Skoog (MS) plates in darkness for 24 h. Interaction was analysed using TCS SP2 (AOBS) laser confocal scanning microscope (Leica Microsystems). Empty vectors pSAT4-DEST-N (1–174 N-YFP) and pSAT5-DEST-C (175-END C-YFP) were bombarded into onion epidermal cells for negative control experiments.

### 2.6 Subcellular localization of At1g08710

F-box gene *At1g08710* was cloned in to pENTD-TOPO vector and transferred to pEarleyGate 104 vector using Gateway cloning technology (Invitrogen, CA). Subcellular localization of At1g08710 was determined using PDS-1000 Helios Gene Gun (Biorad) in onion epidermal cells. These onion peels were kept on Murashige and Skoog (MS) plates in darkness for 24 h. After the incubation, YFP fluorescence was observed using a TCS SP2 (AOBS) laser confocal scanning microscope (Leica Microsystems).

### 2.7 Analysis of mutant seeds of At1g08710

T-DNA insertion mutant seeds of *At1g08710* gene (SAIL_3B04 – T DNA) were obtained from Arabidopsis Biological Resource Centre (https://abrc.osu.edu/). For genotyping genomic DNA was isolated from the mutant plants using DNeasy plant minikit (Qiagen). Homozygous lines of the mutant plants were confirmed by running a PCR reaction on genomic DNA isolated from mutant as well as wild type plant leaves using gene specific and T-DNA left boarder primers (Supplementary table). Further, DNA bands obtained in homozygous plants were excised from agarose gels and sequenced to know the site of insertion. T-DNA was found to be inserted at 1^st^ exonic region of *At1g08710*. Transcript was analyzed for the final confirmation of the mutant seeds.

### 2.8 Seedling growth and stress treatments

All the wild type and mutant plants were grown in plant growth chamber under controlled growth conditions (16/8 h light/dark cycle, temp. 22 ± 2 °C and light intensity 100 μm m^−2^ s^−1^). Seeds were harvested from these plants and kept in the room temperature under dark conditions for two weeks. Seeds from all these genotypes were germinated on normal half-strength MS-agar plates and kept in a growth room under controlled conditions. Seven-day-old seedlings were transferred to half-strength MS-agar media. For Abscisic acid treatment medium was supplemented with 10 10μM ABA. For drought stress, medium was supplemented with Mannitol (150mM). These plates were monitored daily for the observation of stress-induced changes.

### 2.9 Accumulation of H_2_O_2_ and MDA content

Malondialdehyde content was measured according to Heath and Packer (1968). 0.2g of seedling tissue was crushed in 2ml of 0.25% 2-thiobarbituric acid made in 10% (w/v) Trichloroacetic acid. The solution was incubated at 95°C for 30 min and centrifuged at 13000g for 30 min. Supernatant was collected and absorbance was measured at 532nm and at 600nm. Concentration of MDA was calculated using an extinction coefficient of 155 mM^−1^ cm^−1^. Hydrogen peroxide (H_2_O_2_) content was measured according to Alexieva et al. (2001). 0.2g of tissue was crushed in 2ml of 0.1% Trichloroacetic acid (TCA) and centrifuged at 13000 g for 15 min at 4°C. Supernatant was collected and 0.5 ml of supernatant was added to the reaction containing 0.5ml of 10mM potassium phosphate buffer (pH 7.0), 1 ml of 1M potassium iodide (KI) reagent and kept in dark to develop color. Absorbance of the color developed was measured at 390nm. Standard curve was made with known H_2_O_2_ concentrations. The amount of H_2_O_2_ present in the tissue was calculated using standard curve.

## 3. Results

### 3.1. Molecular characterization and subcellular localization of Arabidopsis F-box protein At1g08710

*At1g08710* gene encodes an F-box protein. *At1g08710* cDNA sequence analysis revealed that it contains an open reading frame of 888 base pairs. At1g08710 gene contains four exons and three intronic regions. At1g08710 gene is located on chromosome number one (Fig. 5A). At1g08710 gene encodes a functional protein, which is 295 amino acids in length, it has a molecular weight of 34.34 kDa with an isoelectric point of 9.98. When At1g08710 protein sequence was searched in EMBL-EBI Interproscan database to identify the domains it is found to contain F-box domain in its sequence from 4 to 64 amino acids. The F-box protein also consists coiled-coil domains (95-129/170-190 aa) in its structure (Fig. 1A, Supplementary Fig. S2). Subcellular localization of F-box protein At1g08710 determined through confocal visualization of At1g08710 YFP-fused protein using onion epidermal cells. YFP fluorescence was observed using a TCS SP2 (AOBS) laser confocal scanning microscope (Leica Microsystems). F-box protein was found to be localized to both nucleus and plasma membrane (Fig. 1B).

**Figure 1.**
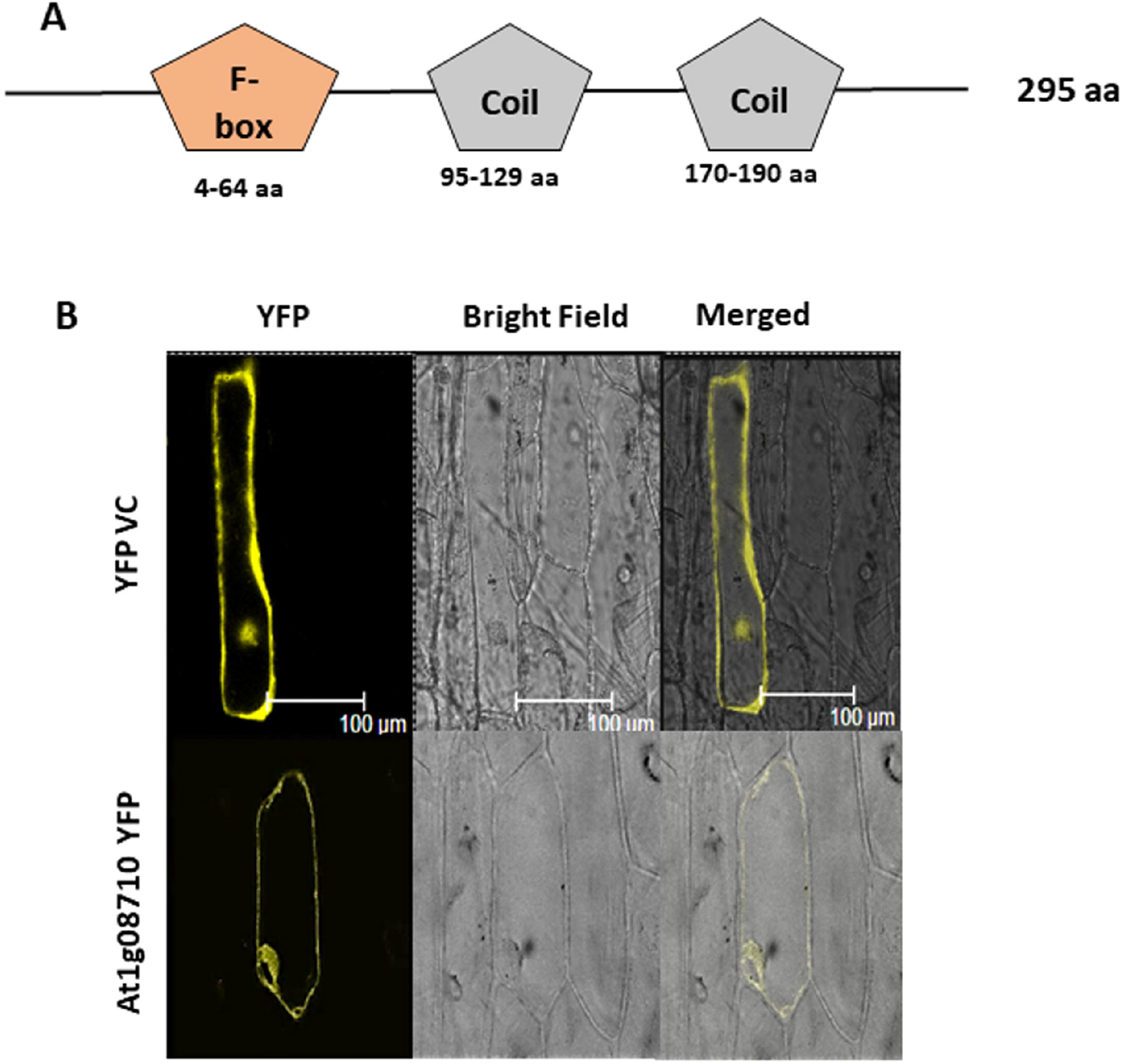
Subcellular localization and domain details of F-box protein At1g08710. (A) F-box protein At1g08710 contains F-box (4-64 aa) and 2 coiled coil domains (95-129 and 170-190 aa) in its sequence as predicted in the protein sequence using EMBL-EBI Interproscan database. (B) F-box protein is localized to both nucleus and cell membrane. F-box protein localization is determined using YFP in onion epidermal cells. YFP empty vector control (VC) is used as a positive control. Bright field and merged images are also shown.

**Figure 2.**
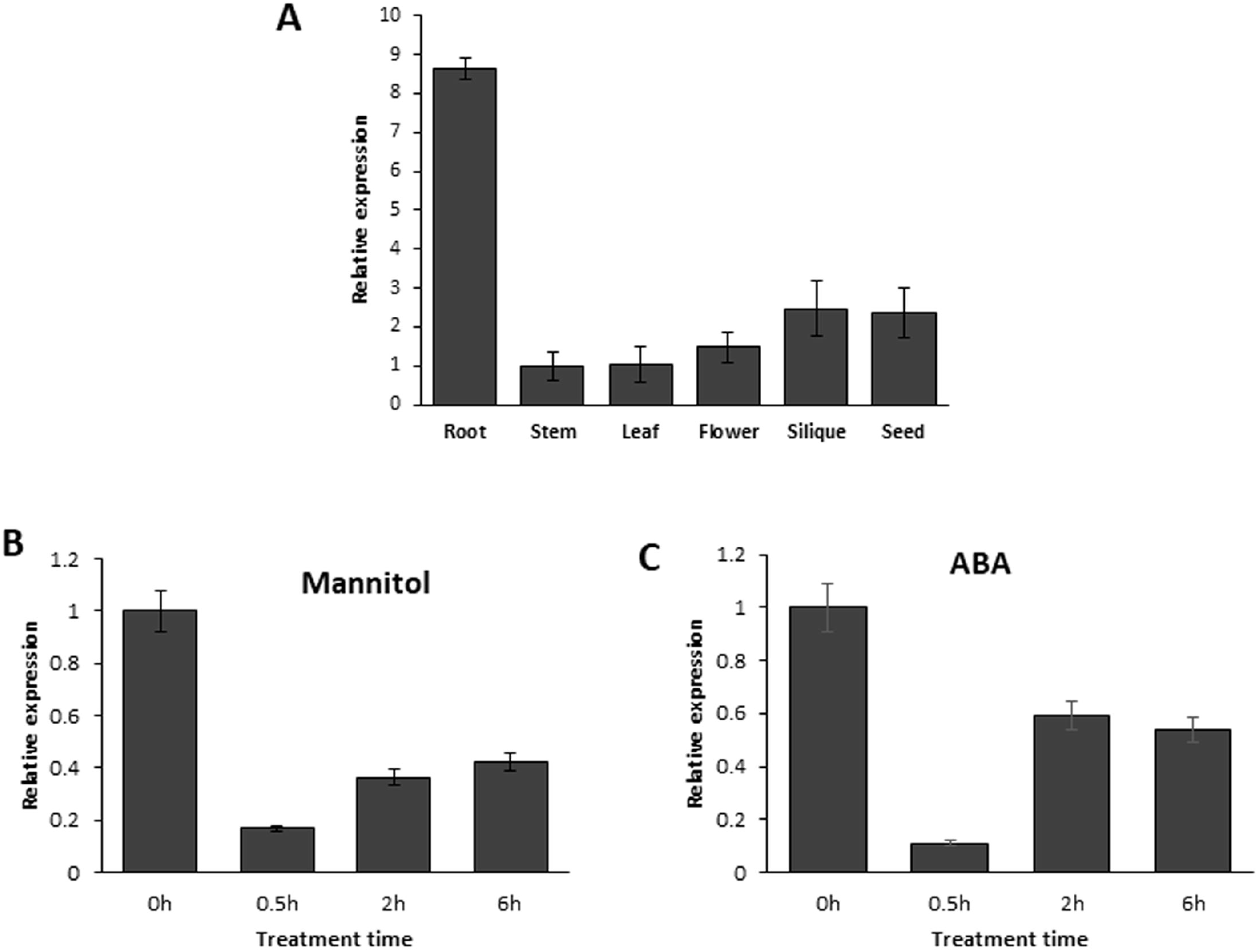
Expression analysis of *At1g08710* in *Arabidopsis* wildtype col-0 plants (A) Quantitative RT-PCR analysis of *At1g08710* in root, stem, leaf, flower, silique and seed tissues of mature plant. (B) Transcript levels of *At1g08710* under drought stress (Mannitol, 150mM) and (C) in the plants treated with Abscisic acid (ABA). 7-day old seedlings were treated with ABA (10μM). Relative expression values were calculated using 2^−ΔΔCT^ method, *18sRNA* is used as an endogenous control. The values are means of three biological replicates and error bars indicate the standard deviation.

**Figure 3.**
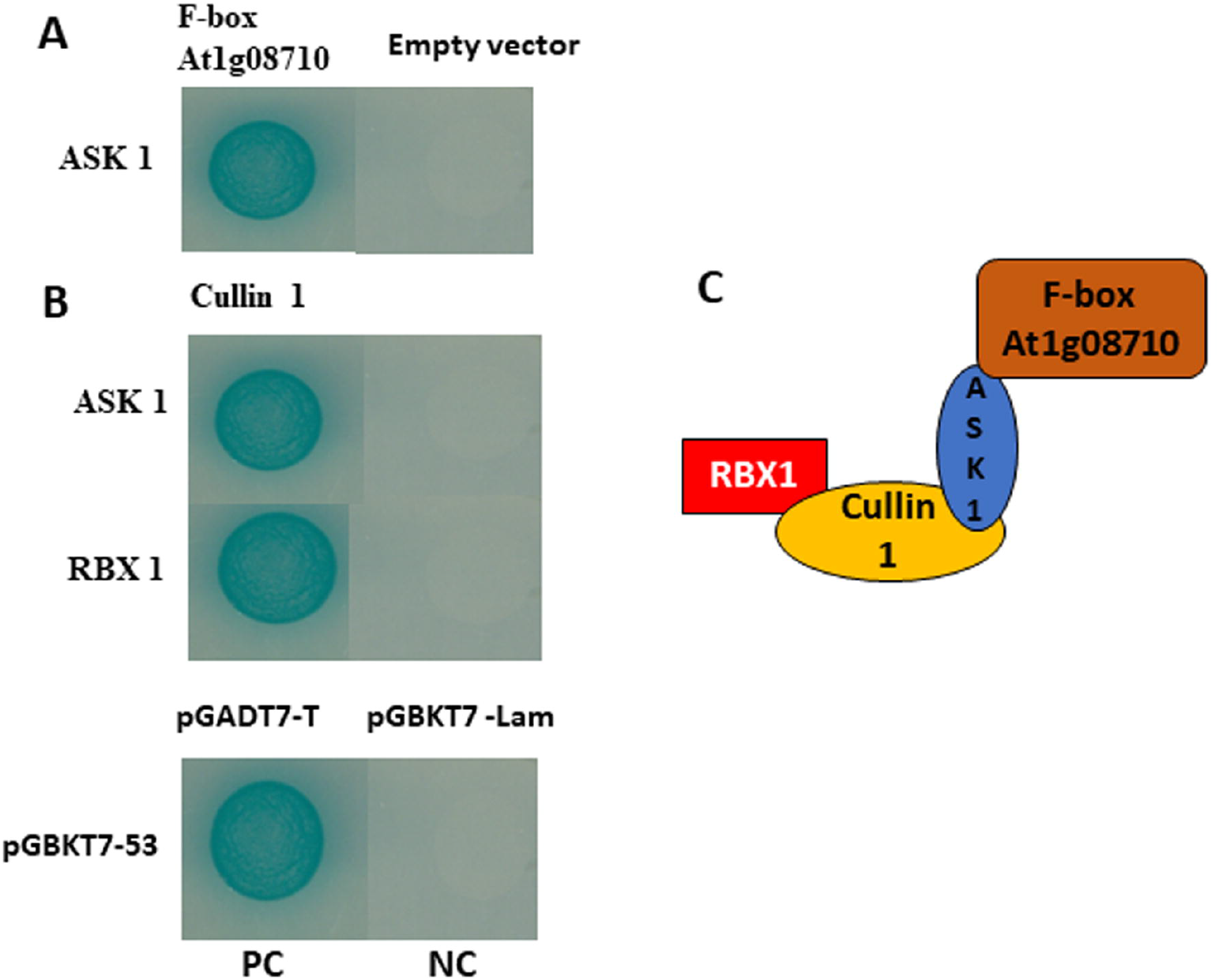
F-box protein At1g08710 interaction analysis with SCF complex subunits. (A) Yeast two-hybrid (Y2H) interaction analysis of F-box protein At1g08710 with ASK1 (*Arabidopsis* SKP1 like protein 1). Yeast cells were co-transformed with pDEST-GBKT7:*At1g08710* and pDEST-GADT7:*ASK1* (B) Y2H interaction analysis of Cullin 1 with RBX1 and ASK1 proteins. Yeast cells were co-transformed with pDEST-GBKT7:*At1g08710*, pDEST-GADT7:*ASK1/RBX1* independently. These cells were grown on quadruple drop-out medium lacking adenine, histidine, tryptophan, leucine and supplemented with X-α-Gal and aureobasidin A. Cells co-transformed with pGADT7-T and pGBKT7-53 were used as positive control (PC). Cells co-transformed with pGADT7-T and pGBKT7-Lam vectors were used as negative control (NC). (C) Schematic diagram illustrating SCF complex subunits.

**Figure 4.**
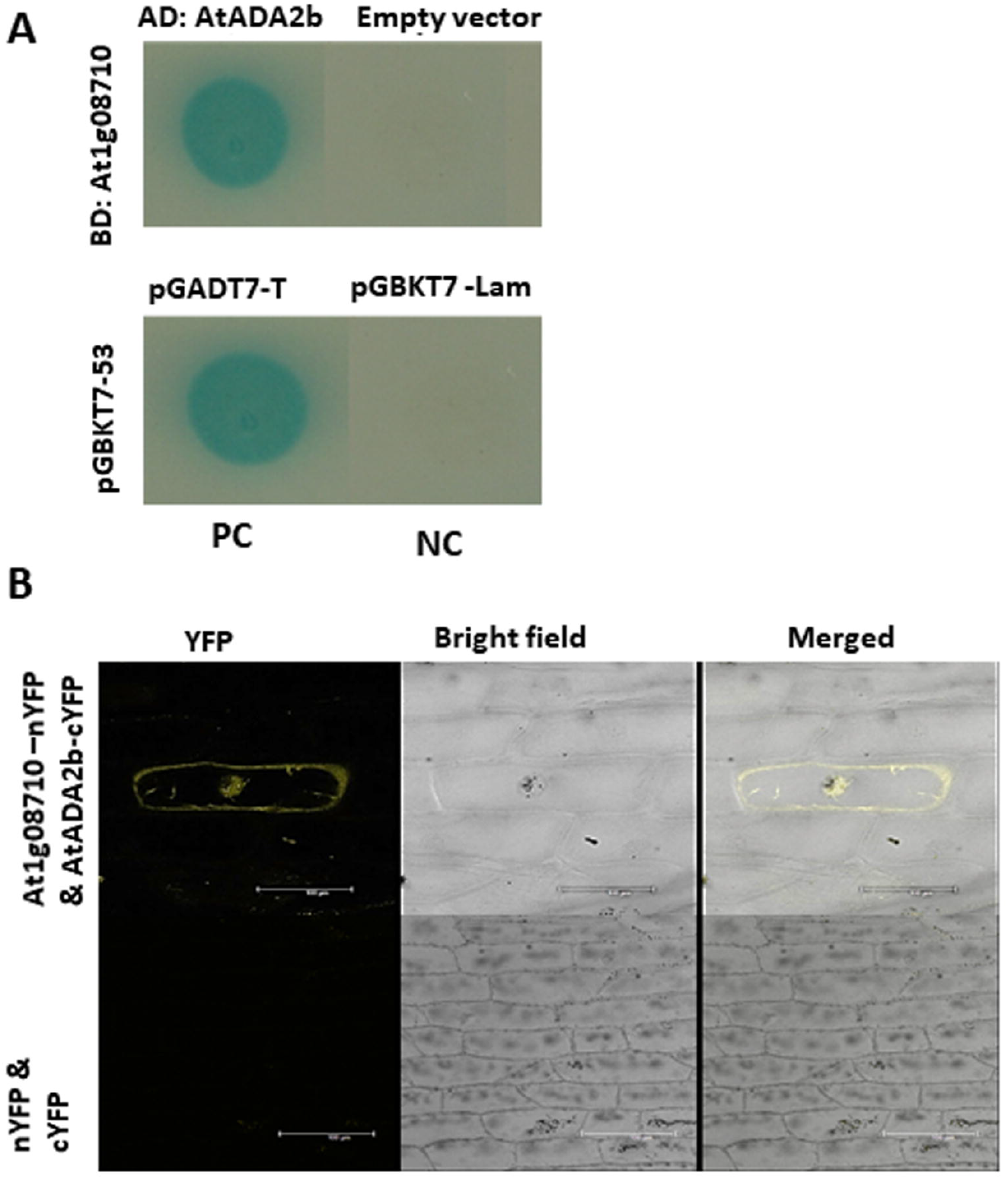
F-box protein At1g08710 interaction analysis with transcriptional coactivator protein ADA2b. (A) Yeast two-hybrid (Y2H) interaction analysis of F-box protein At1g08710 showing its interaction with ADA2b. Yeast cells were co-transformed with pDEST-GBKT7:*At1g08710*, pDEST-GADT7:*ADA2b* and the interaction was analyzed. Cells co-transformed with pGADT7-T; pGBKT7-53 were used as positive control (PC). Cells co-transformed with pGADT7-T; pGBKT7-Lam vectors were used as negative control (NC). (B) Bimolecular fluorescence complementation (BiFC) analysis displaying F-box protein At1g08710 interaction with ADA2b. Onion epidermal cells were co-transfected with pSAT4-DEST-N: *At1g08710*; pSAT5-DEST-C: *ADA2b* and the interaction was analyzed using YFP. Empty BiFC vectors were used as negative control. Bright field and merged images are also shown.

**Figure 5.**
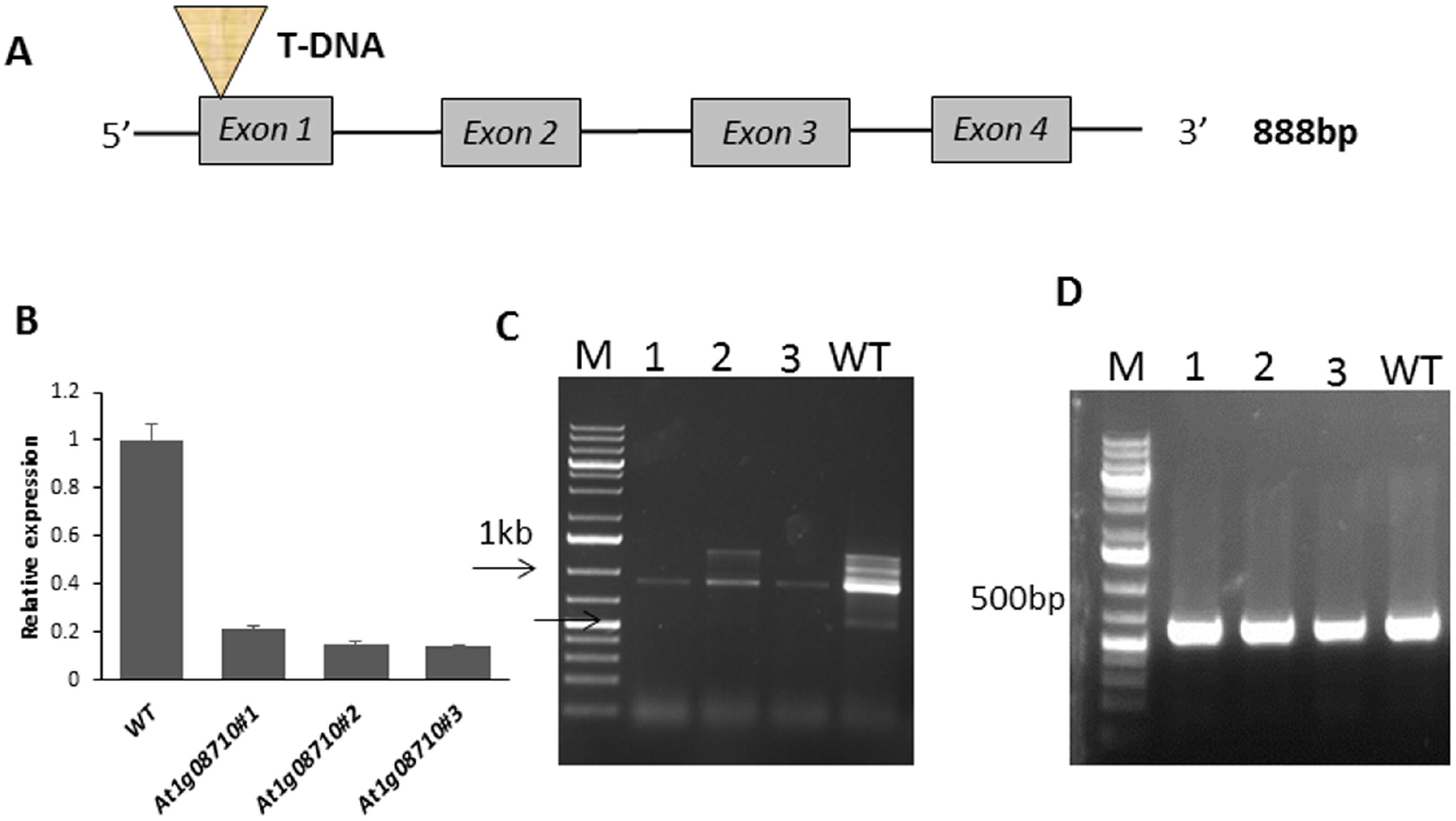
*At1g08710* mutant plants characterization (A) Schematic representation of T-DNA insertion in the F-box gene mutant plants (SAIL_3B04). T-DNA is inserted in the first exonic region. (B) Quantitative RT-PCR analysis of *At1g08710* in the mutant plants. *18sRNA* is used as an endogenous control. (C) Semiquantitave PCR analysis of *At1g08710* in F-box gene mutant plants. (D) Equal quantity of RNA has been used as shown by the *18s RNA* amplification.

### 3.2 F-box gene At1g08710 transcript highly accumulated in root and altered in response to drought stress

To understand the function of Arabidopsis F-box protein At1g08710, initially its transcript was analyzed in different organs of mature plant. To analyze *At1g08710* transcript, RNA was isolated from different organs of mature plant and cDNA was synthesized by reverse transcription PCR. Quantitative real time PCR analysis revealed *At1g08710* is highly expressed in root followed by other organs (Fig. 2A). This result suggests its possible role in root growth. To know whether F-box protein has any role in drought stress, seedlings were exposed to ABA and drought stress conditions. *At1g08710* transcripts was found to be altered in response to ABA and drought stress conditions indicating its role during abiotic stress (Fig. 2B, 2C).

### 3.3 F-box protein At1g08710 interacts with ASK1, Cullin proteins to form SCF complex

SCF E3 ligases are well studied E3 ligases in plants. SCF is a complex of three subunits Arabidopsis SKP1 like protein 1 (ASK1), Cullin and F-box proteins. To verify if F-box protein At1g08710 is part of SCF complex its interaction with SKP1 and Cullin proteins was analysed through yeast two hybrid experiment (Y2H). To perform Y2H, *At1g08710, Cullin 1* were cloned in DNA-binding domain vector pDEST-GBKT7. *ASK1, RBX1* genes were cloned in activation domain vector pDEST-GADT7. Interaction of F-box protein with ASK1 and Cullin 1 interaction with ASK1, RBX1 were analysed independently. These constructs were transformed into Y2H gold strain. Cells which are able to grow and produce blue colour on QDO/X/A medium were considered positive for interaction. Several blue colonies were spotted which revealed the F-box protein At1g08710 interaction with ASK1, and cullin 1 interaction with both RBX1 and ASK1 proteins, thus forming SCF complex (Fig. 3).

### 3.4 F-box protein At1g08710 interacts with transcriptional co-activator protein ADA2b

F-box proteins perform their functions by targeting selective protein for degradation. So in order to elucidate the function of F-box proteins, it is essential to identify their target proteins. To identify the interacting partners of F-box protein At1g08710, Y2H experiment was performed. Y2H experiment was performed according to the manufacturer guidelines using *Arabidopsis* yeast two hybrid library (Clonetech). Transcriptional co-activator protein ALTERATION/DEFICIENCY IN ACTIVATION 2B (ADA2b) is found to be interacting with F-box protein At1g08710 in Y2H assay. To confirm the interaction, Y2H one to one interaction assay was performed. For this F-box gene *AT1G08710* and *ADA2b* were cloned in to pDEST-GBKT7 and pDEST-GADT7 vectors respectively and transformed to Y2H gold strain. F-box protein is found to be interacting with ADA2b (Fig. 4A). To further confirm the interaction, Bi-molecular fluorescence complementation (BiFC) assay was performed. To carry out BiFC, *At1g08710* and *ADA2b* genes were cloned in pSAT4-DEST-N (1–174 N-YFP) and pSAT5-DEST-C (175-END C-YFP) vectors, respectively. Both constructs were introduced in to onion epidermal cells using PDS-1000 Helios Gene Gun (Biorad). In BiFC assay, YFP signal found in nucleus further confirmed F-box protein interaction with transcriptional co-activator protein ADA2b (Bi-FC) (Fig. 4B).

Chromatin undergoes various modifications to regulate the expression of genes. Histone acetylation is one such dynamic epigenetic chromatin modification. Transcriptional co-activator protein ADA2b interacts with histone acetyl transferase GCN5 (Stockinger et al., 2001). Under water deficit conditions, ABA-Responsive Element Binding 1 (AREB1) transcription factor interacts with ABA-Responsive Element (ABRE) promoter. AREB1 further recruits ADA2b-GCN5 complex to induce the expression of various drought responsive genes (Li et al., 2018). Promoter analysis study revealed the presence of drought responsive cis-acting element AtMYB2 BS RD22 in the promoter region of ADA2b gene. ADA2b gene promoter also contains ABA responsive ABI3/VP1 transcription factor binding motif RAV1-A (Supplementary Fig. S4). ADA2b mutant plants have showed pleiotropic phenotypes including aberrant root development, dwarf size, a smaller number of petals and stamens. Root growth is severely affected in mutant plants (Vlachonasios et al., 2003).

Transcript of ADA2b was analyzed in F-box *At1g08710* mutant plants. Under normal conditions, expression of ADA2b is slightly higher in the F-box mutant plants compared to the wild type plants. To check whether it has any role in drought stress adaptation, ADA2b transcript was analyzed in F-box mutant plants under the stress conditions. ADA2b transcript was highly upregulated in the mutant plants under drought stress conditions (Fig.7A). ADA2b transcript was also highly accumulated in mutant plants treated with ABA, suggesting its role in drought stress adaptation (Fig.7E).

### 3.5 Molecular characterization of Arabidopsis At1g08710 mutant plants

Mutant seeds of *At1g08710* (SAIL_3B04 – T DNA insertion lines) were obtained from Arabidopsis biological resource center. To establish genotype, genomic DNA was isolated from both WT and the mutant plants using DNeasy plant minikit (Qiagen). Homozygous plants of the mutant seeds were identified by running a PCR reaction using T-DNA left boarder and gene specific primers. (Supplementary Fig. S3). Further sequencing analysis confirmed the T-DNA insertion in the 1^st^ exonic region of *At1g08710* (Fig. 5A). In order to study the level of transcription of *At1g08710* in the mutant background RNA was isolated and cDNA was made. Transcript levels were found to be knocked down in *At1g08710* mutant homozygous plants (Fig. 5B), Semi quantitative PCR analysis with the cDNA further confirmed the knocked down transcript in the F-box mutant plants (Fig. 5D).

### 3.6 F-box protein At1g08710 negatively regulates root length and imparts drought stress tolerance

Transcript analysis of F-box gene *At1g08710* in various mature plant organs revealed its higher accumulation in roots (Fig. 2A). This prompted us to study F-box protein’s role in root growth. Therefore, we obtained the mutant seeds from ABRC and identified homozygous plants (Supplementary Fig. S3). F-box mutant seedlings displayed longer roots in comparison to the wild type under normal growth conditions suggesting the involvement of F-box protein At1g08710 in controlling root growth (Fig. 6A, 6D).

**Figure 6.**
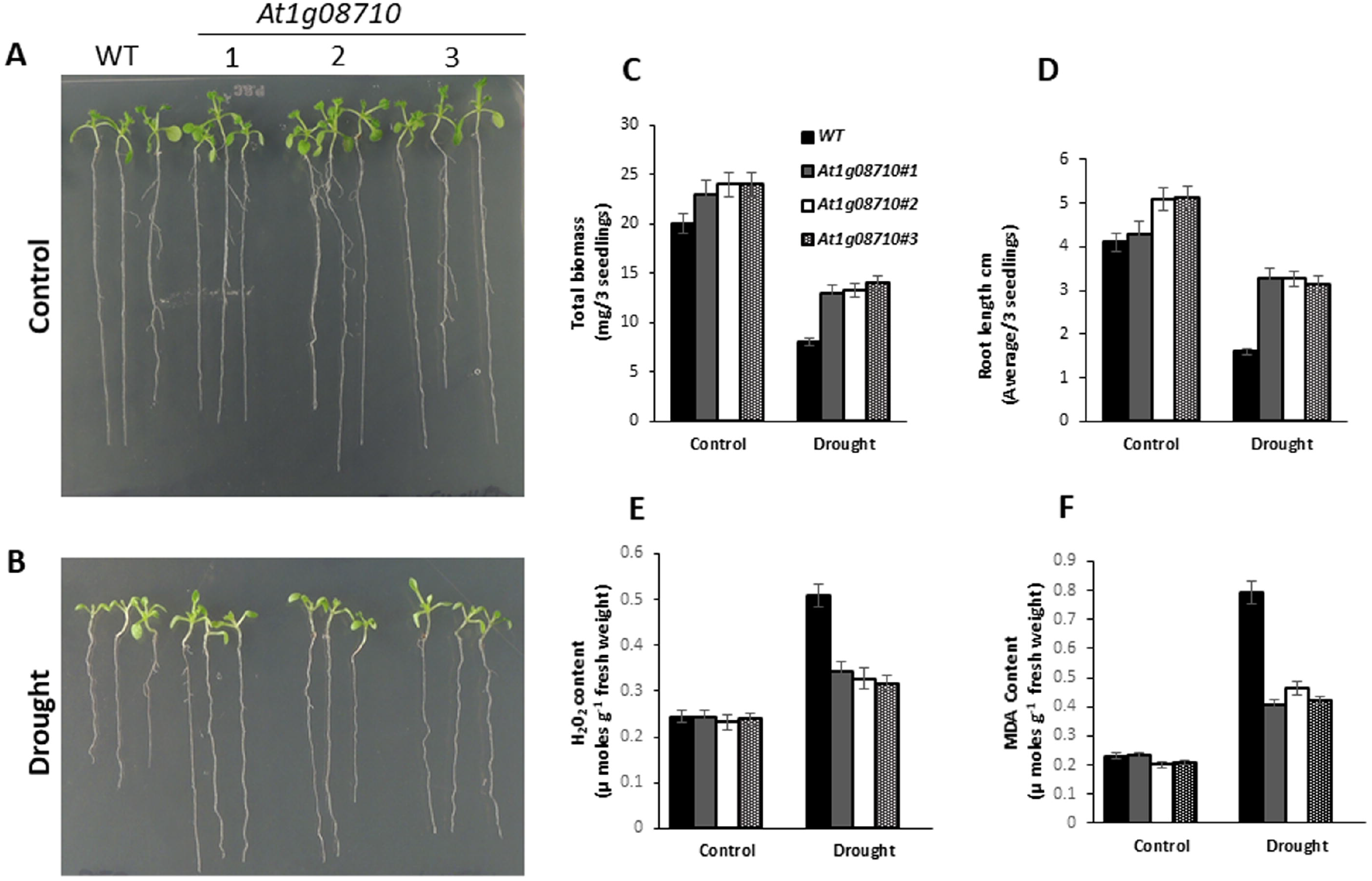
*At1g08710* mutant plants exhibit improved seedling growth under drought stress conditions. (A) comparative phenotypic analysis of wild type (WT), mutant plants under control conditions and (B) under drought stress conditions. Seven-day old seedlings were treated with mannitol (150mM) and the growth pattern was compared. (C) Total biomass of WT, mutant plants under drought stress. (D) Root length of WT, mutant plants treated with mannitol. Quantitative analysis of (E) H2O2 content and (F) MDA content in WT, mutant plants under drought stress conditions. The values are means of three biological replicates and error bars indicate the standard deviation.

Initial transcript analysis revealed *At1g08710* transcript altered during drought stress (Fig. 2B). To investigate whether At1g08710 has any role in seedling growth particularly under drought stress condition, seven-day old seedlings of F-box mutant and wildtype were treated with mannitol (150mM) and growth pattern was observed. *At1g08710* mutant seedlings grew better under drought stress conditions in comparison to wildtype seedlings. *At1g08710* mutant seedlings remained green, healthy and performed better under drought stress conditions (Fig. 6B), whereas wildtype seedlings were severely affected under similar conditions. H_2_O_2_ and MDA contents are normally considered as stress indicators (Verslues et al., 2006; Jambunathan, 2010). When H2O2 and MDA contents were analyzed, F-box mutant seedlings showed significantly reduced H_2_O_2_ and MDA accumulation compared to the wildtype seedlings after stress treatment. These results reveal role of F-box protein At1g08710 in drought stress adaptation (Fig. 6E, 6F).

### 3.7 Drought stress responsive genes expression is induced in F-box gene At1g08710 mutant plants

Environmental stress conditions such as drought adversely affect plant growth and productivity. Plants adopt to drought stress through various molecular, biochemical, physiological and cellular responses. In Arabidopsis, transcript of many genes is induced in response to water deficit conditions. Previous studies have demonstrated the induction of ABA responsive genes such as *AtRD22, AtRD29A, AtERD11, AtDREB1, AtNCED3, AtRAB18, AtCOR47* and *AtABI3* under drought stress conditions (Weng et al., 2014; Min et al., 2015; Liu et al., 2016; Qin et al., 2018; Song et al., 2018). To further understand whether F-box protein At1g08710 has any role in drought stress, transcripts of drought-responsive genes such as *Responsive to Dehydration 29A (RD29A), Responsive to Dehydration 22 (RD22), Abscisic acid insensitive 3 (ABI3)* were analyzed in F-box mutant and wild type plants using qRT-PCR. Under normal conditions expression of *RD29A, RD22, ABI3* is slightly higher in the F-box mutant plants compared to the wildtype plants. Whereas, under water deficit conditions, transcripts of these genes were significantly increased in mutant plants (Fig. 7B, 7C, 7D). These plants were further subjected to ABA treatment and the transcripts of *RD29A, RD22, ABI3* were analyzed. Transcripts of all these genes were highly upregulated in the mutant plants compared to the wildtype plants in response to ABA treatment (Fig. 7F, 7G, 7H). In Y2H, Bi-FC experiments F-box protein was found to be interacting with transcriptional co-activator protein ADA2b. Hence, *ADA2b* transcript was also analyzed under similar conditions in F-box mutant plants to see if it is affected. *ADA2b* transcript was found to be upregulated in plants treated with ABA and water deficit conditions (Fig. 7A, 7E). Taken together, these results reveal mutation in F-box gene leads to the induction of drought responsive genes and thus is involved in drought stress tolerance.

**Figure 7.**
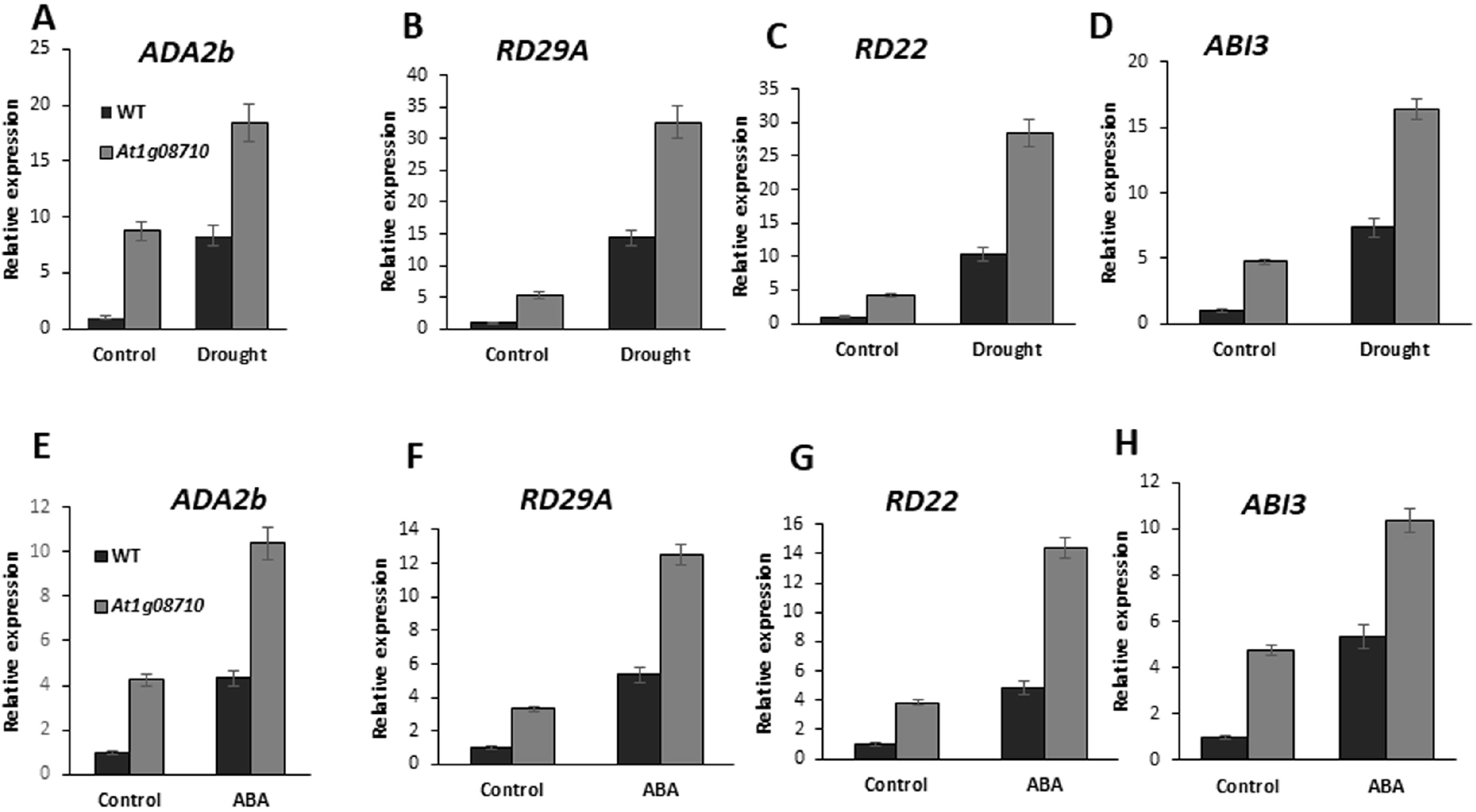
Expression analysis of drought responsive genes in *At1g08710* mutant plants. 7-day old seedlings were treated with (Mannitol, 150mM) and ABA (10μM). Quantitative Real-time PCR analysis of drought responsive *ADA2b* (A, E); *RD29A* (B, F); *RD22*(C, G); *ABI3*(D, H) genes in F-box mutant plants subjected to Mannitol and ABA. Relative expression values were calculated using 2^−ΔΔCT^ method, *18sRNA* is used as an endogenous control. The values are means of three biological replicates and error bars indicate the standard deviation.

## 4. Discussion

Drought stress response in plants is a complex process and is controlled by multiple cellular and molecular signaling events. Recent studies revealed role of ubiquitin mediated 26S proteasome pathway (UPS) in plant response to abiotic stress conditions (Lyzenga et al., 2012). The ubiquitinylation of protein substrate is accomplished by the sequential action of three key enzymes namely ubiquitin activating enzyme (E1), ubiquitin conjugating enzyme (E2) and ubiquitin ligase (E3). Subsequently, the resulting ubiquitin marked protein is recognized and degraded by 26S proteasome complex (Smalle & Vierstra, 2004). Among the three enzymes required for ubiquitination of proteins, ubiquitin ligase E3 is key enzyme that plays crucial role in providing substrate specificity. E3 ubiquitin ligases account for 90% of the UPS genes. SCF E3 ligases is one of the subfamilies of E3 Ubiquitin ligases. F-box protein is an integral member of E3 ligase SCF complex comprising SKP1, Cullin and Rbx1 protein (Gagne et al., 2002). Previous studies reported F-box protein At1g08710 interaction with ASK13 (Rao et al., 2008). To check the ability of F-box protein At1g08710 to form functional SCF complex, F-box protein interaction with ASK1, and Cullin1 protein interaction with both ASK1 and RBX1 was analyzed using yeast two hybrid (Y2H) assay. F-box protein was found to be interacting with ASK1, and Cullin1 was interacting with both ASK1 and RBX1 (Fig. 3) revealing F-box protein forming a functional SCF complex.

F-box proteins are involved in various biological functions by degrading target proteins through ubiquitin proteasome pathway. Recent studies in *Arabidopsis thaliana* revealed F-box proteins role in drought stress adaptation. *Arabidopsis* RCAR3 interacting FLbox protein 1 (RIFP1) is involved in drought stress. *RIFP1* overexpressing plants are sensitive to ABA treatment. Whereas, *rif1* null mutant plants exhibited drought stress tolerance (Li et al., 2016). *Arabidopsis* F-box protein Drought Tolerance Repressor 1 (DOR1) is also involved in drought stress response. Mutant plants of *dor1* are tolerant to drought stress. Whereas, overexpression plants under the similar conditions are sensitive to drought stress (Zhang et al., 2008). Whereas, ABA responsive F-box domain protein 1 (AFBA1) positively regulates drought stress tolerance as AFBA1 overexpression plants are tolerant to drought stress. (Kim et al., 2017). Another *Arabidopsis* F-box protein, more axillary growth 2 (MAX2) has been shown to be involved in drought stress response. Mutant plants of *max2* are hypersensitive to drought stress and ABA treatment (Bu et al., 2014). *Glycine max* F-box protein 176 (GmFBX176) negatively regulates drought stress response as its overexpression plants were severely affected under drought stress conditions (Yu et al., 2020). Similarly, in this study *Arabidopsis* F-box protein At1g08710 is also found to be involved in drought stress response. When tested F-box gene transcript is highly accumulated in root (Fig. 2A) and drought stress conditions alter the gene expression (Fig. 2B, 2C). F-box gene mutant plants exhibited longer roots compared to the wild type plants under normal conditions (Fig. 6A) and displayed better growth compared to the wild type plants under drought stress conditions. Mutant plants have shown longer roots compared to the wild type plants under water deficit conditions they remained green and healthy (Figure. 6B).

Chromatin undergoes structural modifications influencing the expression and regulation of various genes. Histone acetylation is one such chromatin modification. Histone acetyl transferase GCN5 interacts with transcriptional co-activator protein ADA2b (Stockinger et al., 2001). Under drought stress conditions, transcription factor AREB 1 binds to the ABA-Responsive Element (ABRE) element of drought responsive genes. AREB1 recruits the ADA2b-GCN5 protein complex leading to the expression of drought stress responsive genes (Li et al., 2018). In the yeast two hybrid and Bi-FC experiments F-box protein At1g08710 is found interacting with ADA2b (Fig. 4). Subcellular localization studies revealed F-box protein localization to the nucleus further supporting its interaction with ADA2b (Fig. 1B). It is interesting to note that *ADA2b* gene transcript is highly accumulated in water deficit conditions in the F-box mutant plants (Fig. 7A, 7E).

In Arabidopsis, ABA plays a major role in drought stress response. Recent studies have identified many positive and negative regulators of ABA signaling in response to drought stress (Shinozaki and Yamaguchi-Shinozaki, 2007; Hauser et al., 2011). Cis-acting elements such as Drought-responsive element (DRE) and ABA-responsive element (ABRE) are known to be involved in drought response in ABA-dependent manner (Gomez-Porras et al. 2007; Shinozaki et al. 2003). These Cis-acting elements are present in the promoters of many drought responsive genes. Plant hormone ABA biosynthesis is triggered under water deficit conditions, which in turn induces the expression of drought responsive genes such as *RD29 A, RD29 B, COR47, RAB18, KIN1, RD22* and *ABI* genes. Expression of drought responsive genes *RD22, RD29A, ABI3* were analyzed in the *At1g08710* mutant plants under drought stress conditions. Transcripts of all these genes were upregulated under water deficit conditions (Fig.7).

Stress conditions trigger the accumulation of ROS molecules, which leads to the oxidative damage of plant cell organelles (Verslues et al., 2006). To check the levels of drought stress induced accumulation of ROS molecules H_2_O_2_ content was estimated from wildtype and F-box mutant plants under drought stress condition. Excessive levels of H_2_O_2_ is accumulated in wildtype plants compared to the mutant plants under water deficit conditions (Fig. 6E). Malondialdehyde (MDA) content is considered as marker for lipid peroxidation under stress conditions (Jambunathan,2010). To check the lipid peroxidation, MDA content was estimated from wildtype and mutant plants of *At1g08710*. MDA content is seen increased in wildtype plants compared to the mutant plants under drought stress conditions suggesting a key role of F-box protein in drought tolerance (Fig. 6F).

In Summary, At1g08710 encodes a nuclear, membrane localized F-box protein and forms an SCF complex by interacting with SKP1 and Cullin1 proteins. Expression analysis revealed F-box gene is highly expressed in root followed by other organs and the transcript is altered in response to drought stress. Interaction analysis revealed F-box protein’s interaction with transcriptional co-activator protein ADA2b which is known to induce the expression of drought stress responsive genes. ADA2b gene transcript is upregulated in the F-box mutant plants. Subsequently, drought stress responsive genes *RD29A, RD22, ABI3* are also upregulated in the F-box mutant plants under water deficit conditions. F-box mutant plants survived better under water deficit conditions compared to the wildtype plants. F-box mutant plants have shown reduced levels of H_2_O_2_ and MDA content. Taken together, all these results suggest F-box protein At1g08710 plays a role in drought stress tolerance in *Arabidopsis thaliana*.

## Author contributions

VB conceived the study, participated in the design of the study and edited the manuscript. VR designed and conducted the major experiments. VR generated, analyzed data and wrote the manuscript. All authors have read and approved the final manuscript.

## Acknowledgements

This work is supported by Department of Biotechnology (DBT), Government of India. VR thank the DBT-RA programme in Biotechnolgy & Life sciences for the financial support.

