## Supplementary data for "F-box protein At1g08710 negatively regulates root length and imparts drought stress tolerance in *Arabidopsis thaliana*"

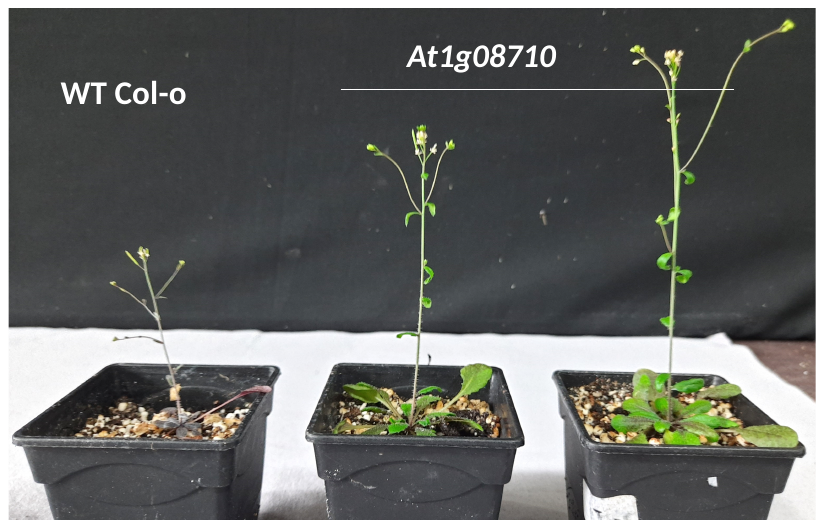

**Supplementary figure S1**. F-box gene *At1g08710* mutant plants are tolerant to drought. Four weeks old wild type and F-box mutant plants were exposed to dehydration stress. Images of plants response after withholding water for 3 weeks followed by 7 days rewatering.

MSANEIPDELWRKILEIGVKSSTFSYKDLCCISISSRRLFRLSCDDSLWDLLLVHDFPNHIVSASSSSESPTKFIYMTRFEREKERKLAAHRRALLRKESEISEWGRRIRELEARLSDEAERLQSSSLQFSDLLKVRQASVALNVWQPEVVRGRQKQMVEQNAVPVEGRLRALEMEMKLCKQQIMGLNRALREVKHRYDIAIKELESMKYHPLRDYKSIRNGDQGSNGKTKKLKTSINYSGDQVSNGKRRKLKTSIDCKFMNISHFSSCSSVTEKFYSYSPKIIHEYIPENLLVL

**Supplementary figure S2.** F-box protein At1g08710 sequence with domain details. F-box gene encodes for a protein consisting 295aa. It contains F-box domain from 4 to 64 amino acids (Highlighted in yellow). The F-box protein also consists of coiled-coil domains in its structure (95-129/170-190 aa - Highlighted in green).

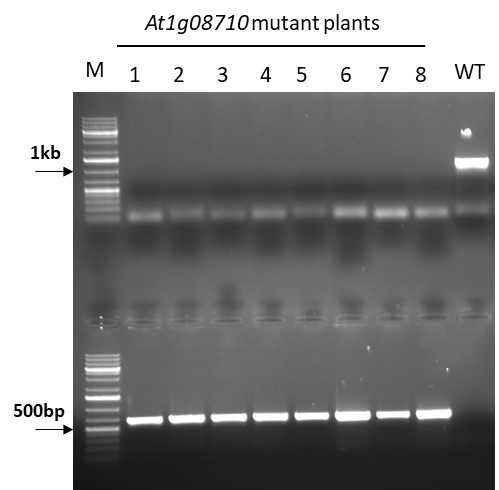

**Supplementary figure S3**. Genotyping of At1g08710 mutant plants. Homozygous lines of the mutant plants were identified by running a PCR reaction on genomic DNA (isolated from F-box mutant, wild type plant leaves) using gene specific primers (upper panel) and T-DNA left border, with gene specific reverse primers (lower panel). Approx. 600bp amplicon specific to T-DNA left border and the gene is seen in plant nos. 1-8 indicating that they are homozygous.

GAGAGAGAGAATATGGGTTTAATCGTAATTTAATACCAAGGACATCATTTCCTTCAGTTTTTTAACTGCAAACTATACTTTTGCCTTTTCTTACATACCAAACCACAAGTCATTGTGGTGTAATATCTATTCAACTACAAGTAAACCCATATCACTGGATTTTTGTCGAGATTATGTGTACATGTTGGGAAATTATTTCGTAGTACAACCTATTATGATGGGAGAATCGAAACTCGTCTTCTGAATTAACTTATCGCAAATCACCGTTAATAAGCGGAAAAGGAATTTACACATCTTACAATTATGCGCAAATACAATTGTTAATTGGATGAGTCATGCCATATTCTTGAGATGGTATGATAGGTATCATTACACCATGTAGTCAATCACGATTTTTTTGGATAATGTGGTTTAGAAGAACTAATCGCGTCTAATATAGACAAACTTATTCGTTCAGCAAAACAACAATTCGATATTTTTATTCGATTGACATGAGTCTTATAAAATTTATCACTAGATGTCTAATAAAAAAGGTTCAAACATTTCTAATAATTCGATTTTCGACCAAAATAAGAAACTTTAAATTAGCATCAAACTATCTTATATGGTTGTTTCCTTGCTTAACCTTCTTCCAAATCAATTTTTCAAATTAGCATCAATAAGATTTTAAAATACATCATAATGTTGTTCCGATAGTGTGTGACAACACTAACCATTTATTCCTATACAAGTTTATTTACTTGCTAATTCATTTACTGAAAATGTCCATGAACAAAATACCTAAAAACGAATAACCATAACCCAACCAAATATTAAAGACATCTCCATCAAGAAGAAACCCAAACAGTTTTTCAAACAAAAAGAATATTAGTATTTTATTATAATTTTATATATGATTAAATTTTTATTTTAGCGAACCAATTGAAAAGAGAAAAATATTATGATGGTTTGTATAAAGTATTTCATAGTTTAACGAGAAACATTCTACACTCTCTTTCTTTTCCTTTTATTATTTTTATTATTTTAATAATAAAAAATTACCAATAAAAATGGTCTATCTTAGCTTTTTTTTCTTCATTTTTTTTTCCCTCAACCACCATTCAACA

**Supplementary figure S4**. Drought responsive Cis-acting elements present in the *Ada2b* promoter (*At4g16420*) region. Promoter analysis revealed the presence of drought responsive cis-acting element AtMYB2 BS RD22 (CTAACCA) in the promoter region of *ADA2b* gene. *ADA2b* gene promoter consists ABA responsive ABI3/VP1 transcription factor binding motif RAV1-A (CAACA). *ADA2b* promoter region also contain other cis acting elements such as MYB4 binding site motif (ACCAAAC) and GATA promoter motifs (TGATAG).

**Supplementary Table: Primers used in this study**

| **Primer No** | **Primer sequence 5’**- **3’** | **Purpose** |
| --- | --- | --- |
| 1 F | CACCATGTCTGCAAACGAGATACCCGACG | *At1g08710* Gateway entry  (F-box gene) |
| 2 R | TCAAAGAACAAGCAGATTTTCAGGAATA |  |
| 3 F | GAAGTTGAAAACAAGCATTGATTGT | *At1g08710* Real time |
| 4 R | GGCATAAAACTCTTCTACATTGCAT |  |
| 5 F | TCCTAGTAAGCGCGAGTCATCA | *18S* rRNA (*At3g41768*)  Real time |
| 6 R | CGAACACTTCACCGGATCAT |  |
| 7 F | CCAGCGATCGTTTATTGCTT | *18S* rRNA Semiquantitative |
| 8 R | AGTCTTTCCTCTGCGACCAG |  |
| 9 F | CACCATGTCTGCGAAGAAGATTGTGTTGA | *ASK1* Gateway entry |
| 10 R | TCATTCAAAAGCCCATTGGTTCTCT |  |
| 11 F | CACCATGGGTCGCTCTCGAGGGAACTTCC | *At4g16420* Gateway entry  (*ADA2b* gene) |
| 12 R | AAGTTGAGCAATACCCTTCTTCACA |  |
| 13 F | CACCATGGAGCGCAAGACTATTGACTTGG | *At4g02570* Gateway entry (*Cullin 1* gene) |
| 14 R | CTAAGCCAAGTACCTAAACATGTTA |  |
| 15 F | CACCATGGCGACTCTAGACTCCGA | At5g20570 Gateway entry  (*RBX 1* gene) |
| 16 R | GTGACCATATTTCTGAAACT |  |
| 17 F | GAACAAGGATGAAAGTGACGG | *At1g08710* genotyping |
| 18 R | TCTCCTTCTCCCTCTCAAACC |  |
| 19 | GCCTTTTCAGAAATGGATAAATAGCCTTGCTTCC | SAIL T-DNA primer |
| 20 F | TCTCCTCCTAAAGTCAAAGT | *ADA2b* Real time |
| 21 R | TACAACAGGTTTCTTTCTAT |  |
| 22 F | CGTTTCTCGGTGGAAGCTGG | *RD22* Real time  *(At5g25610*) |
| 23 R | GCAGTAGAACACCGCGAATG |  |
| 24 F | TAGGTTACTCCGGAGAAATT | *RD29A* Real time  *(At5g52310*) |
| 25 R | TTGTCGTCGTTTCCTTCTTC |  |
| 26 F | AAAAGCAGGATGTATCTCCT | *ABI3* Real time  *(At3g24650*) |
| 27 R | TCATTTAACAGTTTGAGAAG |  |
